# Long-term prevalence data reveals spillover dynamics in a multi-host (*Artemia*), multi-parasite (Microsporidia) community

**DOI:** 10.1101/248542

**Authors:** Eva J.P. Lievens, Nicolas O. Rode, Julie Landes, Adeline Segard, Roula Jabbour-Zahab, Yannis Michalakis, Thomas Lenormand

## Abstract

In the study of multi-host parasites, it is often found that host species contribute asymmetrically to parasite transmission, with cascading effects on parasite dynamics and overall community structure. Yet, identifying which of the host species contribute to parasite transmission and maintenance is a recurring challenge. Here, we approach this issue by taking advantage of natural variation in the community composition of host species. We studied the horizontally transmitted microsporidians *Anostracospora rigaudi* and *Enterocytospora artemiae* in a Southern French metacommunity of their brine shrimp hosts, *Artemia franciscana* and *Artemia parthenogenetica*. Within the metacommunity, patches can contain either or both of the *Artemia* host species, so that long-term prevalence data can provide a direct link between the presence of the two host species and the persistence of the two parasites. First, we show that the microsporidian *A. rigaudi* is a spillover parasite: it was unable to persist in the absence of its maintenance host *A. parthenogenetica*. This result was particularly striking in light of *A. rigaudi’s* high prevalence (in the field) and high infectivity (when tested in the lab) in both hosts. Moreover, *A. parthenogenetica’s* seasonal presence imposed seasonality on the rate of spillover, causing cyclical pseudo-endemics in the spillover host *A. franciscana*. Second, while our prevalence data was sufficient to identify *E. artemiae* as either a spillover or a facultative multi-host parasite, we could not distinguish between the two possibilities. This study supports the importance of studying the community context of multi-host parasites, and demonstrates that in appropriate multi-host systems, sampling across a range of conditions and host communities can lead to clear conclusions about the drivers of parasite persistence.

## Introduction

Although many parasites infect multiple host species within their community (Cleaveland et al. 2001, Taylor et al. 2001), not all hosts are created equal. Host species vary widely in their degree of exposure to the parasite (e.g. Kilpatrick et al. 2006), competence (e.g. LoGiudice et al. 2003, Auld et al. 2017), and population density (e.g. Dobson 1995, Rhodes et al. 1998, Searle et al. 2016). These factors affect their contribution to the parasite’s overall transmission, and thus to the maintenance of a persistent parasite population (Dobson 2004, Streicker et al. 2013).

Quantifying the relative contribution of each host species to the persistence of a multi-host parasite is important for several reasons. Host species that contribute substantially to the parasite’s maintenance have a strong influence on its epidemiology (Viana et al. 2014) and evolutionary trajectory (Holt and Hochberg 2002, Benmayor et al. 2009, Ching et al. 2013), and can be pinpointed in the development of disease control strategies (Fenton and Pedersen 2005, Streicker et al. 2013). Furthermore, asymmetrical transmission of parasites between host species can feed back into community structure (Hatcher et al. 2006).

Unfortunately, identifying which host species contribute to the persistence of a parasite population is notoriously difficult (Viana et al. 2014). Pathogens can persist in a host population if their basic reproduction number R_0_, which represents the number of secondary cases caused by a single infected host, is greater than one (Dobson 2004, Streicker et al. 2013). The host is then said to maintain the parasite population. In multi-host communities, each host species contributes to the parasite’s overall R_0_, and an R_0_ greater than one can mask a lot of variation in those contributions. For example, Fenton et al. (2015) considered a parasite infecting a simple two-host community, finding that it may fall into three underlying categories: *i*) a facultative multi-host parasite, which can be maintained by either host in the absence of the other (R_0_ would still be greater than one if either host was removed from the community); *ii*) an obligate multi-host parasite, which can only persist when both hosts are present (R_0_ would be lower than one if either host was removed); *iii*) a spillover parasite, which can be maintained indefinitely by one of the host species, but not by the other (R_0_ in isolation would be respectively greater and lower than one). In the final case, the parasite’s presence in the second, ‘spillover’ host is dependent on regular reintroductions from the ‘reservoir’ or ‘maintenance’ host (Ashford 1997, Haydon et al. 2002). Distinguishing between these fundamentally distinct, but superficially similar categories is challenging. Lab-based tests of host competence are insufficient, as the epidemiology of the system is key (Searle et al. 2016); solutions therefore include the establishment of epidemiological, statistical, or genetic models, which may be labor-intensive and sensitive to assumptions (Viana et al. 2014).

Another way to identify into which category a multi-host parasite falls is to exploit variation in the composition of natural host communities (Fig. 1). For example, a parasite which can be found in communities containing both hosts *A* and *B*, but also in isolated populations of host *A* or host *B*, is clearly a facultative multi-host parasite. In contrast, a parasite that spills over from host *A* to host *B* can be identified by its presence in communities containing host *A* or hosts *A* and B, but repeated absence in communities containing only host *B*. This type of observation explicitly links the presence or absence of hosts to the presence or absence of disease, and can therefore lead to direct conclusions. Previously, such conclusions have mostly been drawn from human interventions, such as the vaccination or cull of a suspected reservoir (e.g. Dobson 1995, Caley et al. 1999, MacInnes et al. 2001, Nugent 2005, Serrano et al. 2011), or through the experimental construction of host communities (e.g. Power and Mitchell 2004, Searle et al. 2016). However, the approach need not be limited to created variation: we can also look for ‘natural experiments’, host communities whose composition varies in the field. Of course, as natural experiments are not planned, due caution must be taken with regards to potential confounding factors. For instance, the epidemiology of a focal parasite may be shaped by an environmental variable (e.g. temperature, Altizer et al. 2006, Dunn et al. 2006), which happens to covary with the presence of a certain host. To ensure that the conclusions are robust to any such effects, the observations must be repeated across a range of relevant field conditions (e.g. different temperatures).

**Figure 1.**
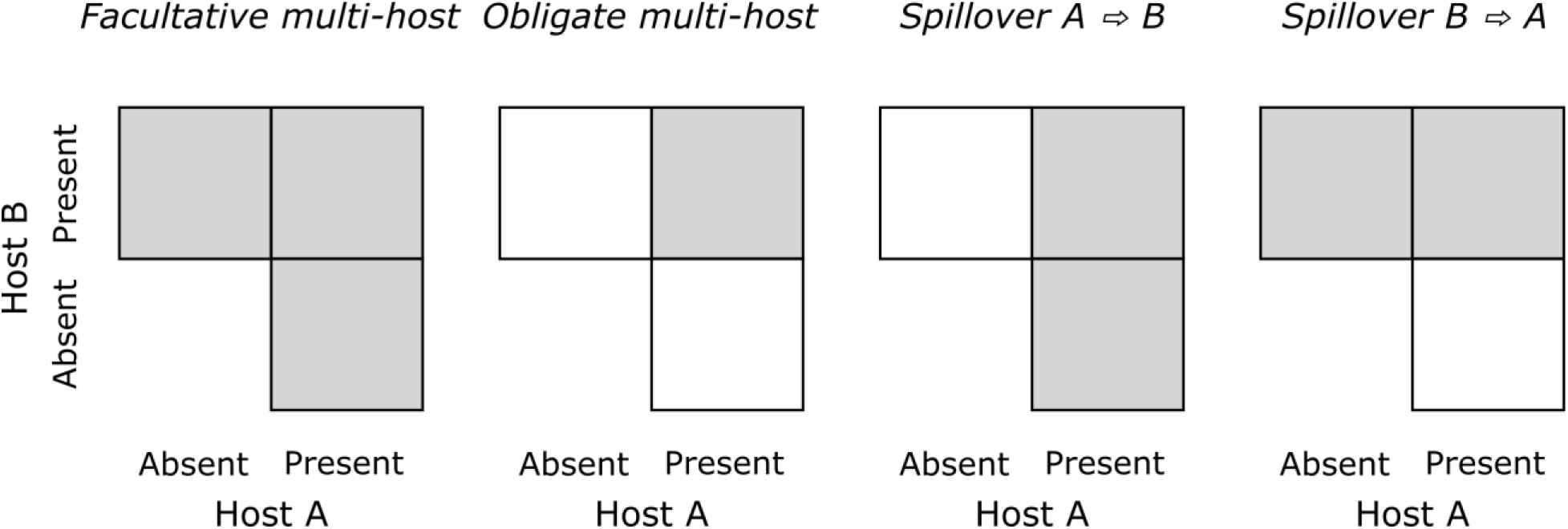
Using variation in the composition of host communities to categorize multi-host parasites. Given similar environmental conditions, the occurrence (gray squares) or absence (white squares) of persistent parasite populations can lead to direct conclusions regarding the parasite’s dependence on its different hosts. Categories following Fenton et al. (2015).

We illustrate the natural experiment approach using a two-host, two-parasite system: the brine shrimp *Artemia franciscana* and *Artemia parthenogenetica*, and their microsporidian parasites *Anostracospora rigaudi* and *Enterocytospora artemiae*. These species occur in sympatry in the French saltern of Aigues-Mortes (Rode et al. 2013a), whose interconnected basins form a metacommunity with patchy species composition. Taking advantage of this variation, we use long-term prevalence data to place the two parasites into the epidemiological categories described above (spillover, facultative multi-host, or obligate multi-host parasites), adding experimental tests of infectivity to investigate some of our conclusions in more depth. We find that the first microsporidian is a spillover parasite, whose spillover dynamics shape its seasonal prevalence. In contrast, we are unable to identify conclusively whether the second microsporidian is a spillover or a facultative multi-host parasite, and we discuss the relative merits of our approach compared to projections based on epidemiological models.

## Methods

### Host-parasite system

*Artemia* (Branchiopoda: Anostraca), also called brine shrimp, is a genus of small crustaceans whose members populate salt lakes and salterns around the world. In Southern France, two *Artemia* species coexist: *A. parthenogenetica* and *A. franciscana. A. parthenogenetica* is a parthenogenetic clade native to the area, while *A. franciscana* is a bisexual species native to the New World (Thiéry and Robert 1992, Amat et al. 2005). *A. franciscana* was first introduced to this region in the 1970’s (Rode et al. 2013c).

*A. rigaudi* and *E. artemiae* are microsporidian parasites of *Artemia* (Rode et al. 2013a). Although highly prevalent in Southern France (Rode et al. 2013c), they have only recently been described and little is known about their ecology. Both species parasitize the gut epithelium and are continuously transmitted to new hosts via free-living spores (Rode et al. 2013b, 2013a). Previously, in a ‘snapshot’ sampling effort of the Mediterranean coast, *A. rigaudi* was found to be more prevalent in *A. parthenogenetica*, while *E. artemiae* was more prevalent in *A. franciscana* (Rode et al. 2013c).

We studied *A. rigaudi* and *E. artemiae* infecting *A. franciscana* and *A. parthenogenetica* in the saltern of Aigues-Mortes, in Southern France. This is a seasonal system, where both temperature (Fig. 2A) and salinity (Fig. 2B) vary throughout the year. *Artemia* hosts are present year-round in large quantities, but their average density varies by more than an order of magnitude between late winter and early summer (estimated at respectively ≤ 1 and 10-15 individuals/L; J. P. Rullmann & P. Grillas, personal communication). The species composition of the *Artemia* community also varies seasonally: *A. parthenogenetica* are entirely absent in winter, but form the majority of the population in summer (Fig. 2C).

**Figure 2.**
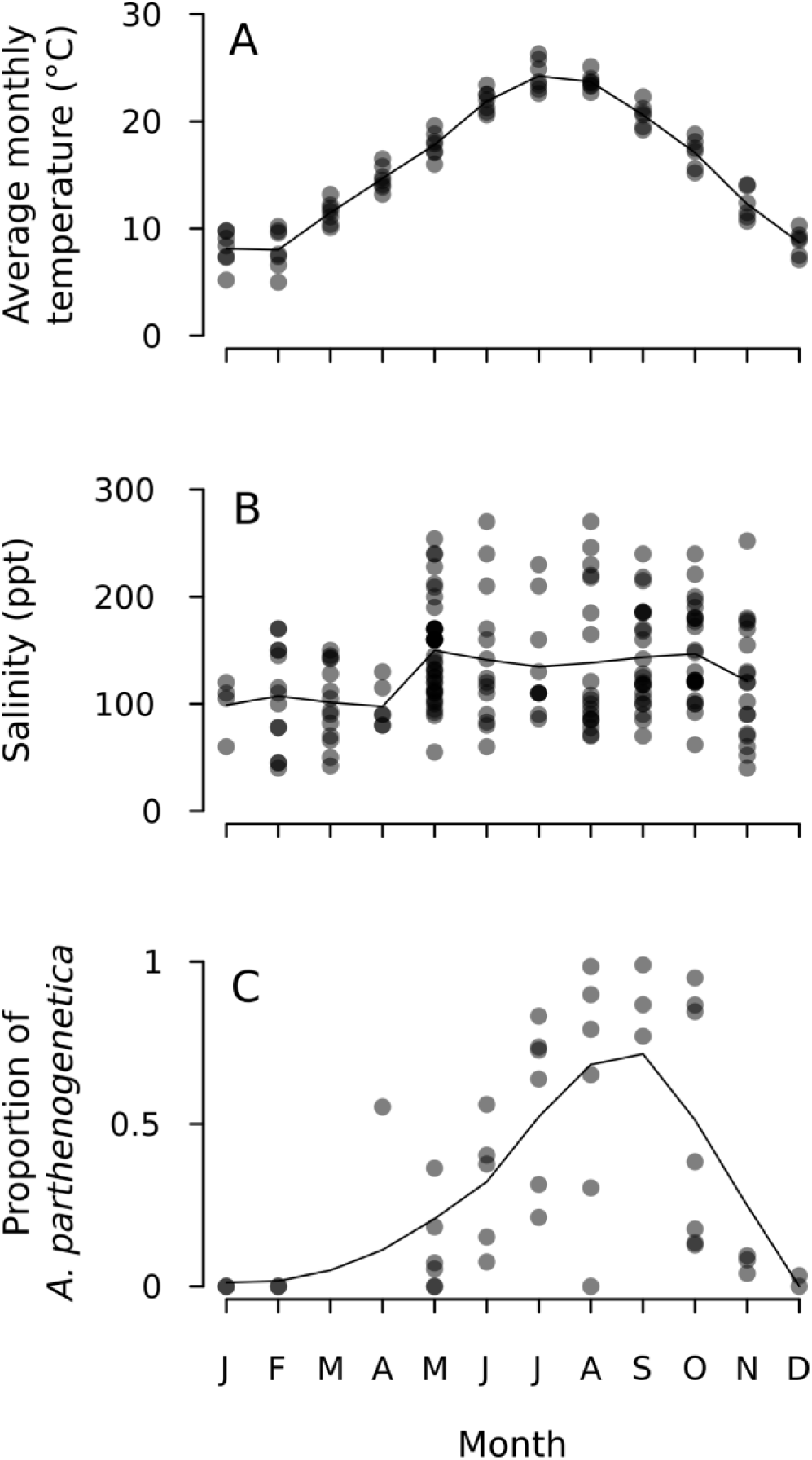
Seasonality in the Aigues-Mortes saltern. Each point represents one data point; overlapping points shade to black. A) Average monthly temperature between 2008 and 2015. Temperature data was collected at the nearby meteorological station Le Grau-du-Roi – Repausset-Levant (Association Infoclimat 2001). The line traces the mean temperature per month. B) Salinity in the Aigues-Mortes saltern, as recorded at various sites between 2008 and 2015 (*n* = 193 observations). The line traces the mean salinity per month. C) Species composition of the *Artemia* community, expressed as the proportion of the population that is *A. parthenogenetica* (figure reprinted from Lievens et al. 2016). All but two of the samples were collected in Aigues-Mortes, the remaining two were collected in Gruissan, France (roughly 100 km South-West of Aigues-Mortes). The line represents a 2^nd^-degree polynomial local regression (LOESS) fitting.

The Aigues-Mortes saltern forms a metacommunity of *Artemia* hosts with patchy species composition. The saltern is made up of a network of large interconnected basins, between which water is allowed to flow or not as a function of the salt production process. This causes environmental factors such as salinity and food quality to vary, leading to variation in the outcomes of inter-host competition: *A. franciscana* or *A. parthenogenetica* can outcompete one another or coexist (Browne 1980, Browne and Halanych 1989, Barata et al. 1996b). At any given time, therefore, adjoining basins can contain different host communities, though gene flow between the basins is regular enough that there is no genetic spatial structure in the *Artemia* populations (Nougué et al. 2015). Within basins, *Artemia* populations are well-mixed (Lenz and Browne 1991), so that we can assume that the microsporidians’ spore pools are shared among the host species (cf. Fels 2006).

### Long-term field data

#### Data collection

We obtained prevalence data for *A. rigaudi* and *E. artemiae* from 94 samples of *Artemia* spp., collected at 14 different sites in the Aigues-Mortes saltern between 2008 and 2015 (Table 1). There was no visible swarming behavior at any of the sampled locations at the time of collection (swarming skews microsporidian prevalence, Rode et al. 2013b). Samples were either processed immediately after collection, or stored in 96% ethanol and processed later; this prevented any infection-specific mortality from skewing the results. We tested a random subset of adult *A. parthenogenetica* and/or *A. franciscana* from each sample for the presence of *A. rigaudi* and *E. artemiae* (mean = 26.8 *Artemia* individuals/sample, *sd* = 24.3). In samples which contained both *A. parthenogenetica* and *A. franciscana*, we usually tested for infection in both host species. Testing was done by PCR using species-specific microsporidian primers, following Rode et al. (2013a).

**Table 1.** Sampling sites in the Aigues-Mortes saltern.

| Site | GPS coordinates | Salinity classification | *n*(samples) |
| --- | --- | --- | --- |
| Caitive Nord | 43° 31' 10.84" N, 4° 14' 26.13" E | middle | 3 |
| Caitive Sud | 43° 31' 10.47" N, 4° 14' 26.12" E | low | 1 |
| Fangouse | 43° 30' 16.04" N, 4° 13' 28.75" E | high | 6 |
| Pont de Gazette | 43° 31' 04.63" N, 4° 10' 48.56" E | low | 15 |
| Puit Romain | 43° 30' 17.83" N, 4° 13' 27.16" E | middle | 6 |
| Site 1 | 43° 29' 53.03" N, 4° 14' 23.06" E | low | 7 |
| Site 3 | 43° 31' 02.65" N, 4° 14' 29.53" E | low | 5 |
| Site 4 | 43° 32' 24.55" N, 4° 13' 25.90" E | middle | 10 |
| Site 5 | 43° 32' 14.96" N, 4° 12' 41.52" E | (unknown) | 1 |
| Site 8 | 43° 31' 37.17" N, 4° 10' 37.77" E | low | 3 |
| Site 9 | 43° 32' 40.31" N, 4° 09' 16.59" E | high | 20 |
| Site 10 | 43° 32' 40.10" N, 4° 09' 17.16" E | high | 3 |
| Site 12 | 43° 31' 55.66" N, 4° 10' 23.92" E | low | 1 |
| St. Louis | 43° 32' 56.78" N, 4° 10' 07.95" E | low | 5 |
| (unknown) | (unknown) | (unknown) | 8 |

In addition to prevalence data, we had environmental and demographic data for most of the samples; we call these variables “sample-specific variables”. First, we knew whether each sample came from a low-, middle- or high-salinity site. The absolute salinity at any given site can change dramatically from one day to the next if the water flow in the saltern is redirected. However, the structure of the saltern means that the salinity at some sites is always lower, higher, or roughly equal to the average salinity of the saltern at that time. We classified these as low-, high-, and middle-salinity sites, respectively (n = 7, 3, and 3 sites; Table 1). This relative classification acts as a residual of salinity after seasonal effects are taken into account, and is not sensitive to the large variability of the absolute salinity measures. We were unable to assign a classification to Site 5, for which we lacked salinity information, or to the 8 samples with an unknown sampling site. Second, we had information on the species composition of each sample: whether it contained both *A. franciscana* and *A. parthenogenetica (n* = 65 samples), only *A. franciscana (n* = 25 samples), or only *A. parthenogenetica* (*n* = 3 samples). One sample had an unknown species composition.

#### Statistical analyses

The goal of our statistical analyses was to identify whether host community composition affected the prevalence of *A. rigaudi* and *E. artemiae*, while controlling for the two major environmental factors in the saltern (temperature and salinity). To do this, we analyzed the prevalence data of each microsporidian species in two steps: first, a general model of the prevalence over time; second, a more complex model that included sample-specific variables. All analyses were performed using generalized additive mixed models (in R version 3.1.3, R Core Team 2015, package “gamm4” Wood and Scheipl 2017), with the number of infected vs. non-infected hosts as the response variable (binomial response with logit link). Model comparison was done using the corrected AIC (Hurvich and Tsai 1989).

First, we constructed models to describe each microsporidian’s yearly prevalence curve, using all of the collected data (n = 138 observations, of which 72 in *A. franciscana* and 66 in *A. parthenogenetica*). These models simply describe the seasonal dynamics of each parasite, i.e. they control for seasonal variation in temperature. We modelled *Prevalence*_ij_ = *Host species*_i_ x s(*Month*_j_) + *Sample*_j_, where *Prevalence*_ij_ represents the proportion of infected host species *i* in sample *j*, s(*Month*_j_) represents a smoothing function of the continuous *Month* variable (the degree of smoothness is adjusted automatically, Wood and Scheipl 2017), and *Sample*_j_ is a random effect controlling for the non-independence of prevalences in *A. franciscana* and *A. parthenogenetica* from the same sample. (We did not include the sampling site in our analyses because working salterns regularly re-distribute the water between basins, so *Artemia* sampled from the same site may or may not have the same genetic background.) The random effect *Sample*_j_ was retained in all of the compared models. The resulting optimal models are hereafter referred to as the ‘general models’.

Second, we investigated whether prevalence was affected by host species composition (our variable of interest) and the sample-specific salinity (which we wish to control for), independently of the variation already explained by our general models. We restricted our dataset to observations of *A. franciscana*, because we had no power to test the effect of species composition on prevalence in *A. parthenogenetica* (the latter was only present by itself in three samples). We further restricted our dataset to sites where the salinity classification and the species composition were known (n = 62 observations in total). For each microsporidian species, we used the general model from the previous section as a null model, and compared it to alternative models with added sample-specific terms. We modelled *Prevalence*_ij_ = (fixed terms of the general model) x (*Presence of* A. parthenogenetica_j_ + *Relative salinity*_j_) + *Sample*_j_, where *Presence of* A. parthenogenetica_j_ and *Relative salinity*_j_ are the sample-specific variables of sample *j*, and *Sample*_j_ is an observation-level random effect accounting for heterogeneity across samples (overdispersion, Harrison 2015). Because *Host species*_i_ was always *A. franciscana*, it was no longer necessary to include this fixed effect, as we did in the general models. The other fixed terms of the general model and the random effect *Sample*_j_ were retained in all of the compared models.

In addition, we tested whether coinfection rates (with both *A. rigaudi* and *E. artemiae*) were significantly higher or lower than expected, as may be the case when parasites inhibit or facilitate each other, or when certain classes of hosts are more or less vulnerable. We used Cochran-Mantel-Haenszel tests (package “stats” in R version 3.1.3, R Core Team 2015) to test the independence of *A. rigaudi* and *E. artemiae* prevalence across samples; these tests were executed separately for *A. franciscana* and *A. parthenogenetica*.

### Experimental tests of microsporidian infectivity

The results of our long-term field data revealed variation in prevalence among host species and on a seasonal basis (see Results). Since field patterns may not reflect functional relationships, we performed two experiments testing the infectivity of *A. rigaudi* and *E. artemiae* in each host and at two temperatures. In both cases, we detected infection using the PCR protocol described by Rode et al. (2013a), and we relied on their finding that an infection is detectable 5-6 days after the host has been exposed to parasite spores.

#### Experiment 1: Effect of temperature and host species/genotype on transmission

We tested the effects of temperature and recipient host species on parasite transmission by exposing non-infected *A. franciscana* or *A. parthenogenetica* to *A. rigaudi* and *E. artemiae* at 15 or 25°C. The purpose of this experiment was firstly to examine the effects of temperature on transmission, and secondarily to provide a rough idea of variation in susceptibility across host species. We did not aim to investigate asymmetric transmission (so we did not add a crossed ‘donor host’ factor), nor to thoroughly document variation in susceptibility across host genotypes (so we used a logistically feasible subset).

The non-infected, ‘recipient’ hosts were adult *Artemia* spp. pulled from laboratory stock collections of Aigues-Mortes lineages. For *A. parthenogenetica*, prior evidence suggests that microsporidian infectivity depends on host genotype (Rode et al. 2013b), so the recipient hosts were collected from four clones, P6 to P9 (P6 and P7 correspond to PAM6 and PAM7 in Nougué et al. 2015). For each clone, we used females from two lines, which had been maintained separately for several generations to standardize maternal effects. We named these recipient groups P6.1, P6.2, P7.1, P7.2, P8.1, P8.2, P9.1, and P9.2. For *A. franciscana*, we formed four replicate recipient groups, F1 to F4.

From each recipient group, 10 individuals were exposed to infection at 15°C, 10 individuals were exposed to infection at 25°C, 5 individuals served as negative controls at 15°C, and 5 individuals served as negative controls at 25°C. The recipient individuals were infected via exposure to infected ‘donor’ hosts. Donor hosts were a mixed group of *A. parthenogenetica* and *A. franciscana*, collected from sites in the Aigues-Mortes saltern with high prevalences of *A. rigaudi* and *E. artemiae* (as ascertained by preliminary PCRs). Using a field-sampled group of mixed donors mimicked natural transmission conditions. Groups of 15 donors were placed in strainers above the jars of 10 recipient hosts. This allowed spores to pass through, but kept donors and recipients from mixing (Rode et al. 2013b). The strainers were rotated every 45 minutes to ensure randomized exposure. Exposure lasted for 9 hours, followed by a 6-day incubation period. On day 6 of the incubation period, the surviving individuals were sacrificed and tested for the presence of *A. rigaudi* and *E. artemiae* as described previously.

#### Experiment 2: Effect of temperature and incubation time on A. rigaudi detection

In experiment 1, both microsporidians had low transmission success at 15°C (see Results). However, the factors underlying this temperature effect were unclear: the reduced transmission could have been caused by a direct effect of temperature on the microsporidians (e.g. on spore germination, Undeen et al. 1993), or by indirect effects of temperature on the ectothermic hosts. Cool temperatures lower the metabolic rate of *Artemia* (Engel and Angelovic 1968), thereby also lowering their defecation and ingestion rates (which would have reduced the effective inoculum size, Burns 1969, Larsen et al. 2008) and dampening their cellular metabolism (which could have slowed the accumulation of microsporidian DNA in the host, Dunn et al. 2006). Since *E. artemiae* was present in the field in winter (see Results), we could infer that its low transmission success at 15°C must be due, at least in part, to indirect effects on the hosts. However, for *A. rigaudi*, which was not found in the field in winter (see Results), it was important to disentangle these confounding effects. To do this, we designed an experiment to compare the effects of temperature and incubation time on (detected) infectivity, while maintaining a standardized spore dose. We limited this experiment to testing *A. rigaudi* infecting *A. franciscana*.

In this experiment, we allowed *A. rigaudi* infections to incubate at 15°C or 25°C for different lengths of time (6 days vs. 12 days). We exposed adult *A. franciscana* from an uninfected laboratory stock population to feces containing *A. rigaudi* spores. The feces were collected from a laboratory stock of *Artemia* spp. infected with *A. rigaudi*; the spore concentration was unknown. Exposure occurred in groups: six groups of 20 hosts were placed in 50 mL autoclaved brine and 2.8 mL fecal solution was added. Four groups were exposed at 15°C, while two groups were exposed at 25°C. Exposure lasted two days, during which time all spores could be ingested (Reeve 1963). After two days, hosts were separated and each individual was placed in a hemolymph tube containing 2.5 mL brine; this prevented between-recipient infections later in the experiment. The infection was allowed to incubate at the exposure temperature for four days, after which half of the surviving hosts from each group were sacrificed, and two of the groups exposed at 15°C were moved to 25°C. After a further six days of incubation, all remaining hosts were sacrificed and tested for the presence of *A. rigaudi*.

#### Statistical analyses

Statistical analyses of the experiments were performed using generalized linear mixed models (package “lme4”, Bates et al. 2015, in R version 3.1.3, R Core Team 2015), with the number of infected vs. non-infected hosts as the response variable (binomial response with logit link). The significance of the predictors was tested using likelihood ratio tests.

For Experiment 1, we analyzed the probability of detecting an infection separately for *A. rigaudi* and *E. artemiae*. Fixed effects included *Temperature*, a *Species/Genotype* factor, and their interaction. As we expected to find differences among the *A. parthenogenetica* clones (Rode et al. 2013b), but could not distinguish between the mixed-together *A. franciscana* families, the *Species/Genotype* factor comprised 5 levels: *A. franciscana, A. parthenogenetica* P6, *A. parthenogenetica* P7, *A. parthenogenetica* P8, and *A. parthenogenetica* P9. We included *Recipient group* as a random effect to control for shared genetic and environmental effects.

For experiment 2, our statistical analyses used *Exposure temperature, Periods incubating at 15°C*, and *Periods incubating at 25°C* as fixed effects (with one period equal to six days), with *Host group* as a random variable controlling for pseudo-replication. Since the sensitivity of our PCR was fixed, an increase in detectability over time reflects an increase in the quantity of parasite DNA present in the host (i.e. intra-host parasite reproduction).

## Results

### Long-term field data

First, we described the prevalence dynamics for *A. rigaudi* and *E. artemiae* throughout the year (Fig. 3, Supp. Table 1). *Anostracospora rigaudi* was strongly seasonal: it was highly prevalent from August to October, but absent in winter *(Month* effect, ΔAICc ≥ 52.7; Fig. 3A & B). These seasonal dynamics were not different in the two hosts, but its prevalence was higher in *A. parthenogenetica* (effect of *Host species*, ΔAICc ≥ 156.2; Fig. 3A vs. B). The prevalence of *E. artemiae* was highly variable and this microsporidian was not strongly seasonal; nevertheless there was statistical support for temporal effects. The precise dynamics depended on the host species: *E. artemiae*’s prevalence increased more steeply towards the end of the year in *A. parthenogenetica* than in *A. franciscana* (interaction between *Host species* and *Month* effect, ΔAICc ≥ 4.8; Fig. 3C & D). Overall, *E. artemiae* was more prevalent in *A. franciscana* (Fig. 3D vs. C).

**Figure 3.**
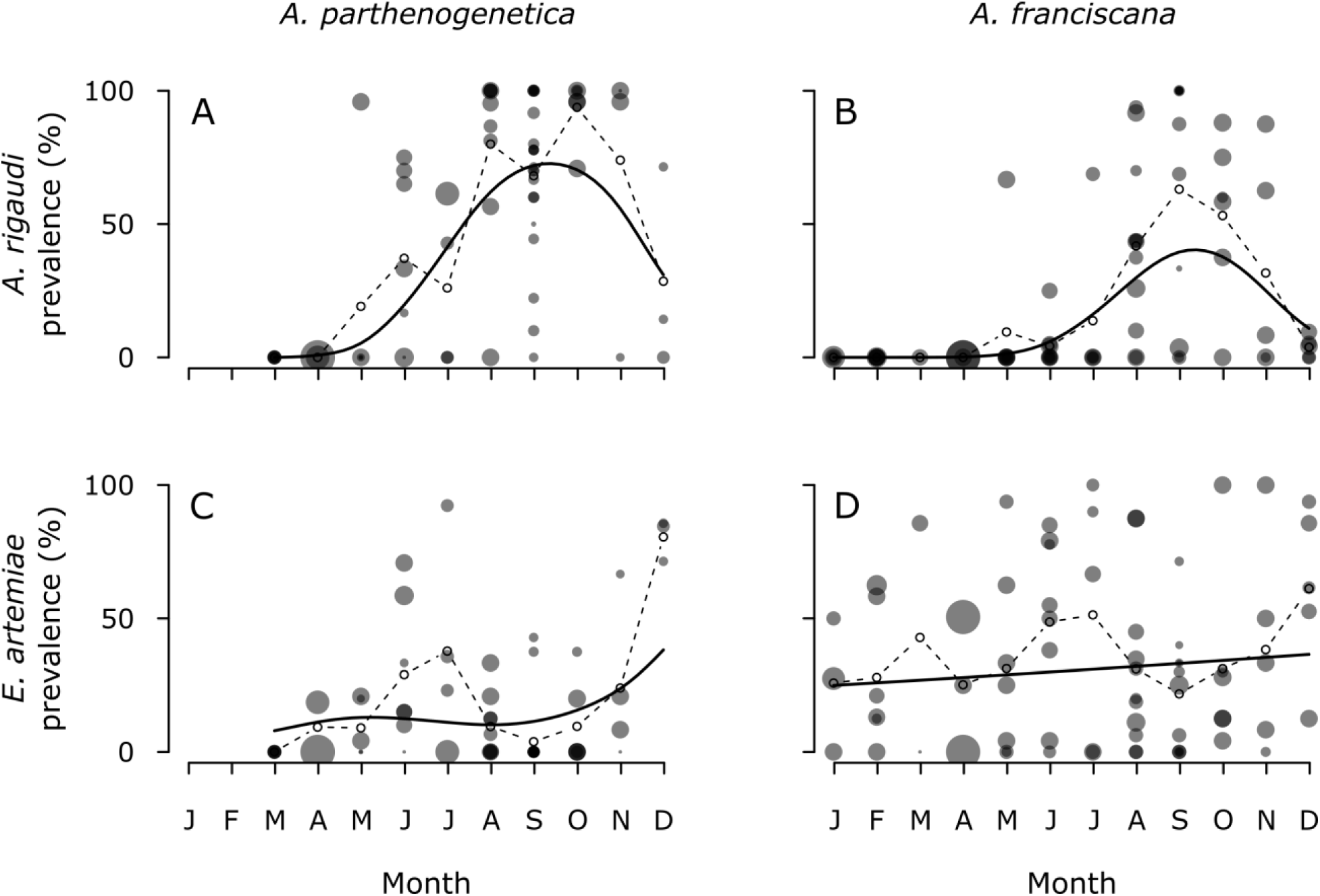
General prevalence patterns of *A. rigaudi* (A & B) and *E. artemiae* (C & D) parasites in *A. parthenogenetica* (A & C) and *A. franciscana* (B & D) hosts. Solid dots are samples; the area of the dot represents the number of individuals in the sample. Overlapping dots shade to black. Solid line: predictions of the general model, represented by the marginal mean (obtained here by averaging over the predictions for all random effects). Using the marginal mean was necessary to compensate for the high variability between samples. Dashed line with open circles: mean prevalence across samples. There is no prevalence data for *A. parthenogenetica* in January and February because this host is absent in winter.

Next, we introduced sample-specific effects to the general models, investigating the effects of salinity and the presence of *A. parthenogenetica* on the prevalence of the microsporidians in *A. franciscana* (Fig. 4, Supp. Table 2). For *A. rigaudi*, the species composition had a strong effect ΔAICc ≥ 5.0): in the absence of *A. parthenogenetica*, the prevalence of *A. rigaudi* was almost always 0%, and never higher than 10% (Fig. 4B). In contrast, in the presence of *A. parthenogenetica* the prevalence could be very high, and the seasonal dynamics found in the general model reappeared (Fig. 4A). There was good support for an additional effect of salinity classification (ΔAICc = 2.1), with the high-salinity sites typically having lower prevalences. For *E. artemiae*, the salinity classification had no effect, and there was little support for an effect of species composition (ΔAICc ≤ −2.1).

**Figure 4.**
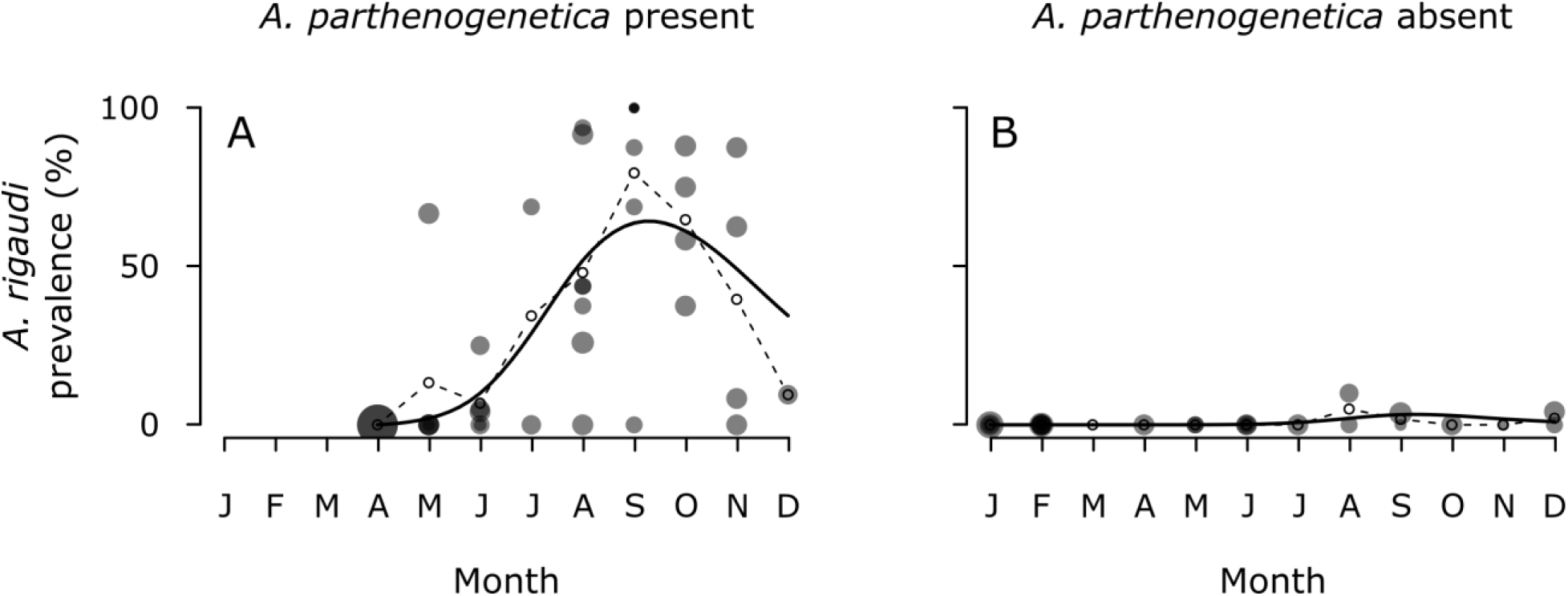
Decomposition of Fig. 2B: prevalence of *A. rigaudi* in *A. franciscana* when *A. parthenogenetica* is present (A) or absent (B). Solid dots are samples; the area of the dot represents the number of individuals in the sample. Overlapping dots shade to black. Solid line: predictions of the best sample-specific model, represented by the marginal mean (obtained here by averaging over the predictions for all random effects). Using the marginal mean was necessary to compensate for the high variability between samples. Dashed line with open circles: mean prevalence across samples.

Finally, coinfection rates varied from 0% to 83% in our samples. Infection with *A. rigaudi* and *E. artemiae* was independent for *A. parthenogenetica* (Mantel-Haenszel χ^2^(1) < 0.1, *p* = 0.83, common odds ratio = 0.8), but was positively associated for *A. franciscana* (slightly more coinfection observed than expected; Mantel-Haenszel χ^2^(1) = 5.1, *p* = 0.02, common odds ratio = 1.9).

### Experimental tests of microsporidian infectivity

#### Experiment 1: Effect of temperature and host/genotype on transmission

The P8 genotype of *A. parthenogenetica* and the F4 replicate of *A. franciscana* were previously infected (all controls tested positive for *A. rigaudi* and *E. artemiae*, respectively), so these recipient groups were removed from the analysis. Surprisingly, the previous infections appeared to have an inhibitory effect: none of the exposed P8s were infected with *E. artemiae* at the end of the experiment (n = 38), and none of the exposed F4s were infected with *A. rigaudi (n* = 21).

Temperature had a clear effect on the probability of infection of both *A. rigaudi* and *E. artemiae*; neither infected well at low temperatures (χ^2^(1) = 136.3 and 46.5, *p* < 0.001 in both cases; Fig.5). *Anostracospora rigaudi* infected all three genotypes of *A. parthenogenetica* and *A. franciscana* equally well (χ^2^(3) = 0.8, *p* = 0.85), but infectivity was dependent on *Species/Genotype* in *E. artemiae* (χ^2^(3) = 18.1, *p* < 0.001; Fig. 5B). Post-hoc Tukey tests indicated that *A. franciscana* and P9 were equally susceptible to *E. artemiae*, while P6 and P7 were much less susceptible (*z* ≤ −2.9, *p* < 0.01). There were no significant interaction effects between temperature and *Species/Genotype*.

**Figure 5.**
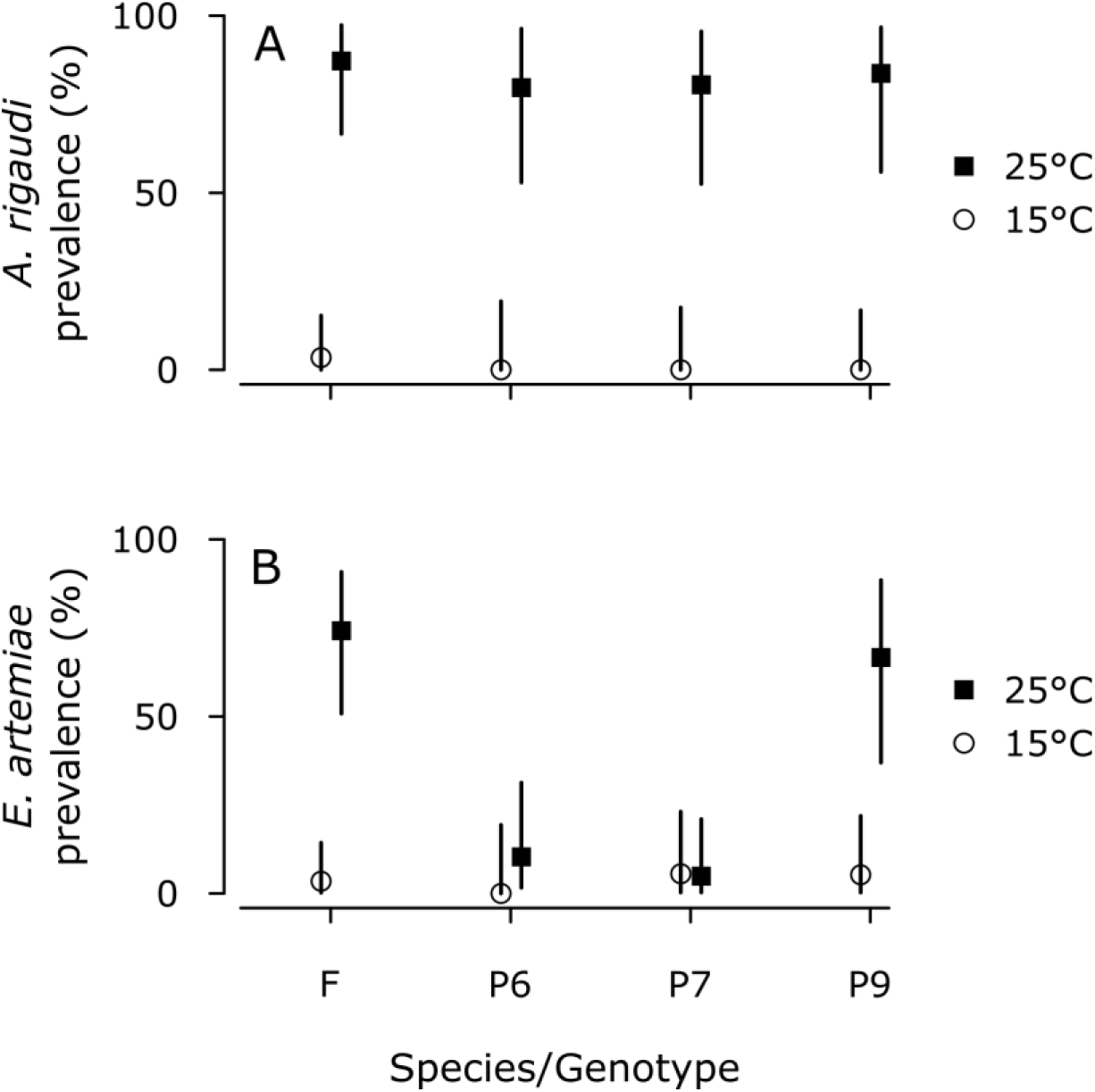
Infectivity of *A. rigaudi* (A) and *E. artemiae* (B) as a function of temperature and host type (experiment 1). Individuals were sacrificed and tested after six days of exposure and incubation at 15°C or 25°C. Species/Genotypes were *A. franciscana* (F) and *A. parthenogenetica* isofemale lines P6, P7 and P9. Vertical lines represent the 95% confidence intervals.

#### Experiment 2: Effect of temperature and incubation time on A. rigaudi detection

The probability of detecting *A. rigaudi* in *A. franciscana* – i.e. the within-host accumulation of parasite DNA – increased significantly with incubation at 25°C, but not at 15°C (χ^2^(1) = 31.9 and 1.9, *p* < 0.001 and *p* = 0.18, respectively; Fig. 6). There was no significant effect of the exposure temperature on infectivity (χ^2^(1) = 0.0, *p* = 0.91). Therefore, the apparent reduction in infectivity of *A. rigaudi* at 15°C during experiment 1 was at least partly caused by slower parasite reproduction inside the hosts, delaying its detectability. The lower rates of detection after 6 days of incubation at 25°C in this experiment compared to experiment 1 could be explained by a lower initial spore dose.

**Figure 6.**
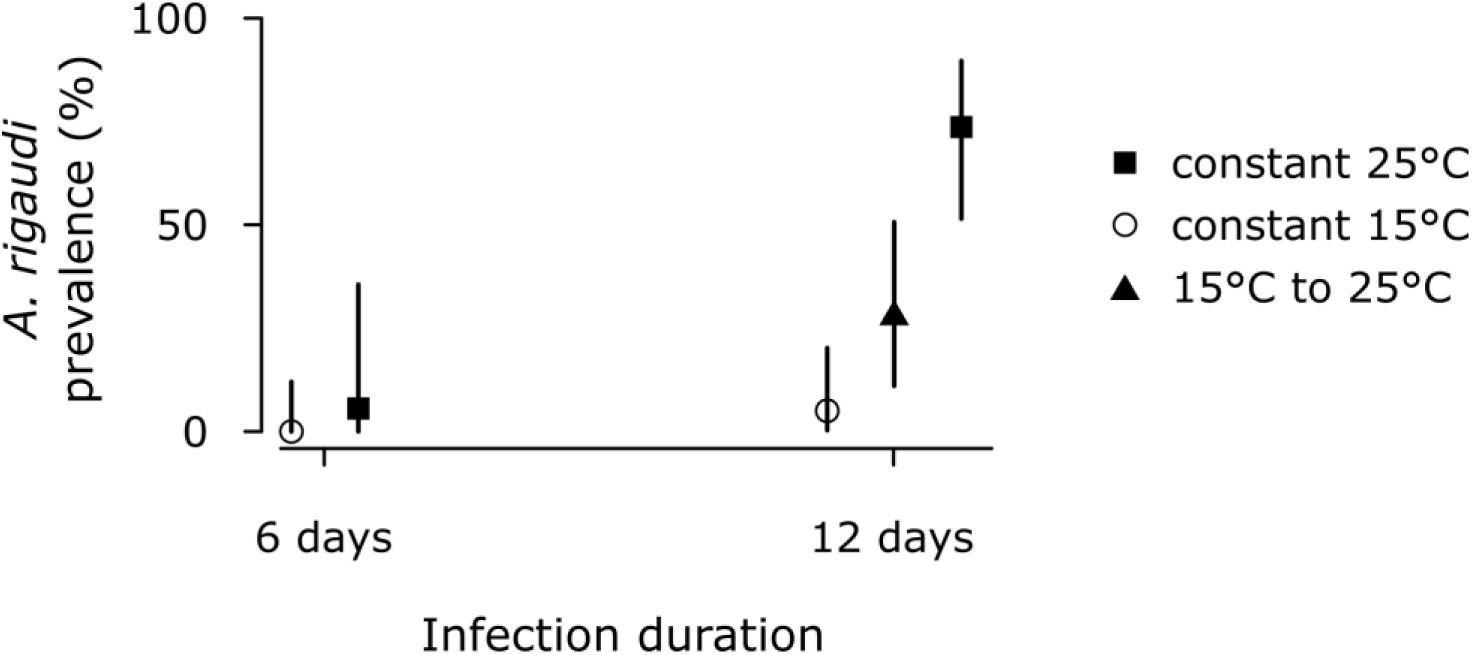
Detection of *A. rigaudi* infections in *A. franciscana* as a function of temperature (experiment 2). Groups of individuals were exposed to *A. rigaudi* and maintained at 15°C or 25°C for six or twelve days, after which they were sacrificed and tested. Two groups were exposed and maintained at 15°C for the first six days, and then moved to 25°C for the remaining six days. Vertical lines represent the 95% confidence intervals.

## Discussion

When a parasite infects multiple host species, we cannot gain an adequate understanding of its epidemiology and evolution without knowing which of the host species actually contributes to its transmission and maintenance. In this study, we exploit natural variation in the composition of *Artemia* host communities to identify their microsporidian parasites as either spillover, facultative multi-host, or obligate multi-host parasites (Fenton et al. 2015). Our primary result is that *A. rigaudi* is a spillover parasite, whose high infectivity in both hosts causes pseudo-endemic dynamics. Secondarily, we show that *E. artemiae* may be a spillover or a facultative multi-host parasite, and speculate that it is the first. Finally, we reflect on the link between the microsporidians’ host specificity and seasonal dynamics, and discuss the merits of our method.

### A. rigaudi *is (secretly) a spillover parasite*

Our long-term prevalence data revealed that *A. rigaudi* is a spillover parasite: it cannot persist on *A. franciscana* in the absence of *A. parthenogenetica* (Fig. 4). The presence of the parasite in *A. franciscana* must therefore be dependent on regular re-introductions from *A. parthenogenetica*, its maintenance host. This result is robust to variation in temperature (the effect can be found in every season, Fig. 3) and salinity (the effect is found across all salinity categories, results not shown), which are the main environmental factors that affect this system. Under the relevant natural conditions, populations of *A. franciscana* are unable to maintain *A. rigaudi*.

Parasites are unable to persist in a host species if that host provides little or no transmission, which may occur if its susceptibility, abundance, and/or lifetime spore production are low (Dobson 2004, Streicker et al. 2013). Our experiments ruled out the first possibility for *A. franciscana* and *A. rigaudi*, as both hosts were equally susceptible to the microsporidian (Fig. 5A). The second possibility, that sites containing only *A. franciscana* have consistently lower host densities, we also consider to be unlikely. Although the dataset used here does not contain demographic information, both observation and available evidence show that *A. franciscana-* only sites can reach similar densities as sites containing both hosts (J. P. Rullmann & P. Grillas, unpublished data). Instead, *Artemia* biomass is mainly constrained by food availability (Browne 1980), temperature (Barata et al. 1996a), and salinity (Wear and Haslett 1986), which are seasonal environmental factors (see Fig. 1). We therefore predict that the third possibility is most probable, namely that *A. franciscana* is a poor host for *A. rigaudi* because infected hosts produce few spores. Further experimental studies will be needed to confirm this hypothesis.

The contrast between *A. rigaudi’s* generalist infectivity and status as a spillover parasite is particularly interesting. At first glance, its high infectivity in the two hosts, as tested in the lab and reflected in the field prevalences (Figs. 4A, 2A & B), may tempt us to conclude that *A. rigaudi* is a generalist parasite. Only a more detailed analysis of the field data belies this generalist infectivity, and reveals *A. rigaudi’s* ‘secret’ host specificity. This result is a good demonstration of the dangers of using only infectivity as an indicator of parasite specialization and long-term success (Agosta et al. 2010, Lange et al. 2015). In addition, the high infectivity of *A. rigaudi* in *A. franciscana* means that spillovers from *A. parthenogenetica* to *A. franciscana* occur frequently enough that the parasite appears to be independently present in both hosts (Fig. 2A & B). Some authors have termed such frequent spillovers ‘apparent multi-host’ or ‘pseudo-endemic’ dynamics (Fenton and Pedersen 2005, Viana et al. 2014)(cf. Dobson 1995, Rhodes et al. 1998), terms which explicitly indicate the difficulty of distinguishing such dynamics from true endemism. This makes the danger of misinterpreting the epidemiology and evolution of pseudo-endemic parasites very high, if their community context is not investigated. *Anostracospora rigaudi* itself is a good example of this problem. Based on previously available information, an earlier paper proposed that *A. rigaudi* could overwinter by infecting the invasive *A. franciscana*, thereby increasing its negative impact on the population of native *A. parthenogenetica* (a process known as “spillback”, Rode et al. 2013c). In contrast, our current conclusion suggests that *A. franciscana* individuals act as inhibitory hosts, absorbing more *A. rigaudi* spores than they produce (Holt et al. 2003), and thereby diluting the effect of *A. rigaudi* on *A. parthenogenetica* (Ostfeld and Keesing 2000, Hall et al. 2009).

### E. artemiae *may be a spillover or a facultative multi-host parasite*

Our prevalence data identified *E. artemiae* as either a spillover or a facultative multi-host parasite, but was insufficiently powerful to distinguish between the two possibilities. We were unable to analyze the ability of *E. artemiae* to persist on *A. parthenogenetica* because only three of our samples did not contain *A. franciscana*. Although these three samples were *E. artemiae-* free, this may have been due to chance. Based on our experimental test of infectivity, however, we suspect that *E. artemiae* is also a spillover parasite, in this case from *A. franciscana* to *A. parthenogenetica*. At high temperatures, *E. artemiae* is very infective in *A. franciscana* and the *A. parthenogenetica* genotype P9, but not at all in the genotypes P6 and P7 (Fig. 5B). These findings are consistent with the genotype-dependent prevalence data collected earlier by Rode et al. (2013b) in natural populations. Since *A. parthenogenetica* populations are a mix of different genotypes (Nougué et al. 2015), we can expect *E. artemiae* to infect *A. parthenogenetica* less well on average. (This interpretation is consistent with our field observations, but some caution is still merited. Since we only tested a small number of genotypes, it is possible that our results would change given a more thorough evaluation of the within-host genetic variation for resistance (cf. Luijckx et al. 2014).) The spillover effect of *E. artemiae* might therefore be even stronger than that of *A. rigaudi*, as the former appears to be more specifically infective than the latter. In the absence of conclusive evidence, however, this microsporidian serves as an excellent example of the difficulty of using observational data to identifying host contributions to parasite success (see discussion below).

### Presence of maintenance hosts shapes parasite seasonality

When comparing the seasonal prevalence patterns of *A. rigaudi* and *E. artemiae* (Fig. 2) with the seasonal changes in host community composition (Fig. 1C), it quickly becomes clear that the two are linked. *Anostracospora rigaudi’s* maintenance host, *A. parthenogenetica*, is entirely seasonal, being completely absent in winter and highly prevalent in late summer and autumn; logically, therefore, *A. rigaudi* is also strongly seasonal. In contrast, *E. artemiae* is able to persist throughout the year, with no overarching seasonal prevalence pattern. By testing infectivity at warm and cool temperature, and comparing the two parasites, we can gain further insight into *A. rigaudi’s* absence in winter. Although low temperatures did slow parasite reproduction within the host when tested in *A. rigaudi* (Fig. 5), they did not make infection impossible for either parasite *(A. rigaudi:* filled triangle in Fig. 5; *E. artemiae:* open circles in Fig. 4B), and clearly do not preclude persistence of *E. artemiae* (Fig. 2D). Therefore, it is possible that *A. rigaudi* could persist in winter, if given the opportunity to do so by the presence of *A. parthenogenetica*. We therefore conclude that the general prevalence patterns of *A. rigaudi* and *E. artemiae* are predominantly shaped by host specialization, and not by environmental factors.

The seasonal cycles of *A. rigaudi* in its spillover host *A. franciscana* are of particular interest (Fig. 2B). These are clearly caused by the seasonal presence of the maintenance host *A. parthenogenetica*, which enforces seasonality on *A. rigaudi* and therefore on the frequency of spillovers. Seasonal cycles in prevalence occur in many host-parasite systems, and are often explained by factors such as climate, host behavior, and host immunity (Hosseini et al. 2004, Duffy et al. 2005, 2009, Grassly et al. 2005, Altizer et al. 2006, Lass and Ebert 2006). In contrast, our study presents a rare example of seasonal infections within a focal host (*A. rigaudi* in *A. franciscana*) being caused by the seasonality of an entirely different host *(A. parthenogenetica*). We know of one other description of such spillover-driven seasonality. Amman et al. (2012) investigated the prevalence of Marburg virus in bats, which causes severe hemorrhagic disease when it spills over to humans. They found that viral infections in bats peaked during birthing seasons, and that 83% of spillovers to humans occurred during these peaks. Such cases highlight the interconnected nature of communities hosting multi-host parasites, and are obviously of particular interest for the establishment of control strategies.

### Extrapolating host contributions from observational data

In this study, we used variation in the composition of natural host communities to investigate how each host contributes to the maintenance of a shared parasite. The strength of this approach is illustrated by the results for *A. rigaudi:* we obtained direct evidence that under natural conditions, it is a spillover parasite dependent on *A. parthenogenetica* and unable to persist in *A. franciscana*. This conclusion, which we could not have drawn from tests of infectivity alone, has crucial consequences for our interpretation of the dynamics and evolution of both parasite and host. On the other hand, the results for *E. artemiae* highlight an important weakness of the method, namely that all possible combinations of communities must be sampled in order to obtain conclusive evidence. If one combination does not occur in the field, this constraint is inescapable. In this case, methods based on the construction of full epidemiological models, such as those described by Fenton et al. (2015), may be more useful. These methods first quantify each host’s contribution to parasite transmission using observed data on host abundance, parasite prevalence, and parasite shedding; they then infer the consequences for persistence from these results. While these techniques require the accurate measurement of all relevant parameters and an adequate mathematical description of the epidemiology, they are not dependent on wide-scale sampling across communities. Such methods are therefore useful where large-scale sampling is difficult or community composition is invariable, while the natural experiment approach could be particularly suited to highly variable and regularly sampled multi-host systems (e.g. *Daphnia*, Ebert 2008). Independently of the method used, these (and our) studies consistently demonstrate the importance of studying a parasite within its entire host community before making inferences about its host specificity, epidemiological drivers, and selection environment.

## Acknowledgements

The authors thank F. Gout, O. Nougué, E. Flaven, and L.-M. Chevin for help with sampling, E. Flaven for help with testing of sampled *Artemia*, and E. Decaestecker, M. A. Duffy and C. L. Murall for their constructive comments on a previous version of the manuscript. We also thank the Salins d’Aigues-Mortes for access to the saltern. We acknowledge support from the QuantEvol (206734) grant from the European Research Council.

## Appendices

**Supplementary Table 1.**
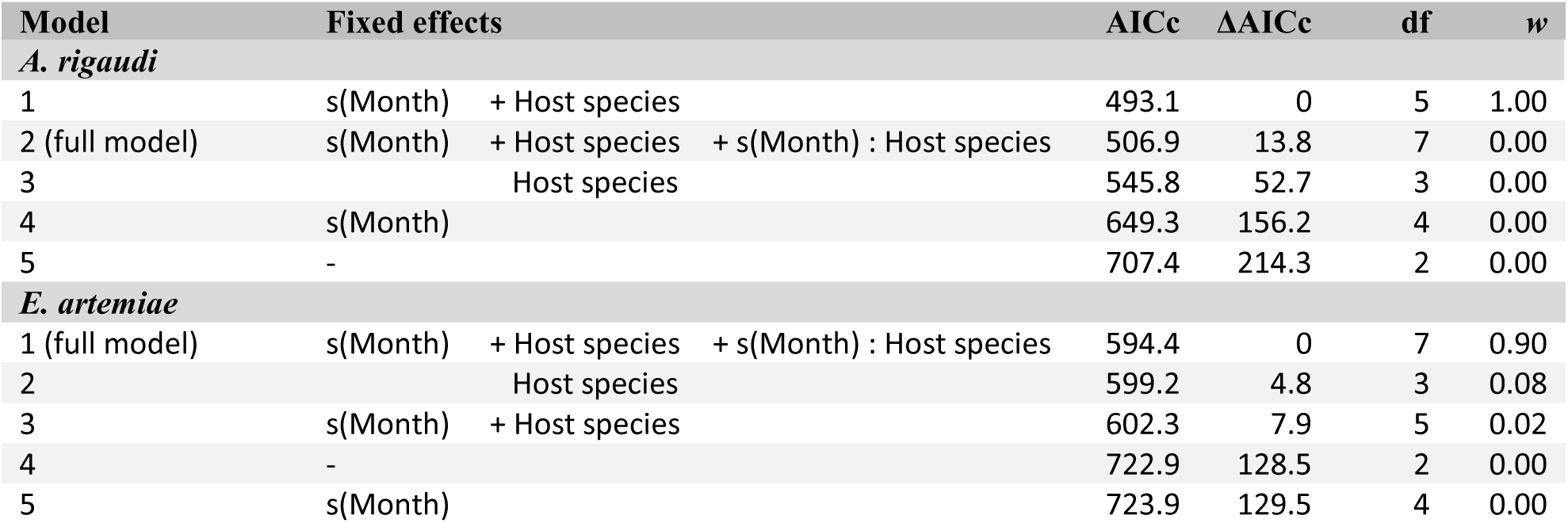
Results of the overall models for *A. rigaudi* and *E. artemiae* infecting both *Artemia* hosts, before the addition of sample-specific variables. The full models included a smoothing function of *Month, Host species*, and their interaction. All models also included *Sample* as an individual-level random effect, to control for pseudoreplication. Models were compared using the difference in corrected AIC(ΔAICc); also provided are the degrees of freedom used (df) and the Akaike weights (*w*).

| Model | Fixed effects |  |  | AICc | ΔAICc | df | w |
| --- | --- | --- | --- | --- | --- | --- | --- |
| *A. rigaudi* |  |  |  |  |  |  |  |
| 1 | s(Month) | + Host species |  | 493.1 | 0 | 5 | 1.00 |
| 2 (full model) | s(Month) | + Host species | + s(Month) : Host species | 506.9 | 13.8 | 7 | 0.00 |
| 3 |  | Host species |  | 545.8 | 52.7 | 3 | 0.00 |
| 4 | s(Month) |  |  | 649.3 | 156.2 | 4 | 0.00 |
| 5 | - |  |  | 707.4 | 214.3 | 2 | 0.00 |
| *E. artemiae* |  |  |  |  |  |  |  |
| 1 (full model) | s(Month) | + Host species | + s(Month) : Host species | 594.4 | 0 | 7 | 0.90 |
| 2 |  | Host species |  | 599.2 | 4.8 | 3 | 0.08 |
| 3 | s(Month) | + Host species |  | 602.3 | 7.9 | 5 | 0.02 |
| 4 | - |  |  | 722.9 | 128.5 | 2 | 0.00 |
| 5 | s(Month) |  |  | 723.9 | 129.5 | 4 | 0.00 |

**Supplementary Table 2.** Results of the sample-specific models for *A. rigaudi* and *E. artemiae* infecting *A. franciscana*. The full models included the fixed factors of the general model (only *s(Month); Host species* could not be included because we only used *A. franciscana*), *Presence of* A. parthenogenetica, *Relative salinity*, and their interactions with the smoothing function of *Month*. All models also included *Sample* as an individual-level random effect, to control for overdispersion. Models were compared using the difference in corrected AIC(ΔAICc); also provided are the degrees of freedom used (df) and the the Akaike weights (*w*). Apart from the general mode, only the models that fell within a cut-off value of ΔAICc = 4 are shown.

| Model | Fixed effects | AICc | $\Delta\text{AICc}$ | df | $w$ |
| --- | --- | --- | --- | --- | --- |
| ***A. rigaudi*** |  |  |  |  |  |
| 1 | *s(Month)* + Presence of *A. p.* + Relative salinity | 204.6 | 0 | 7 | 0.61 |
| 2 | *s(Month)* + Presence of *A. p.* | 206.7 | 2.1 | 5 | 0.21 |
| ... | ... | ... | ... | ... | ... |
| 15 (general model) | *s(Month)* | 224.5 | 20.0 | 4 | 0.00 |
| ... | ... | ... | ... | ... | ... |
| ***E. artemiae*** |  |  |  |  |  |
| 1 (general model) | *s(Month)* | 347.7 | 0 | 4 | 0.64 |
| 2 | *s(Month)* + Presence of *A. p.* | 349.7 | 2.1 | 5 | 0.23 |
| ... | ... | ... | ... | ... | ... |

